# Efficient masking of plant genomes by combining kmer counting and curated repeats

**DOI:** 10.1101/2021.03.22.436504

**Authors:** Bruno Contreras-Moreira, Carla V Filippi, Guy Naamati, Carlos García Girón, James E Allen, Paul Flicek

**Affiliations:** European Molecular Biology Laboratory, European Bioinformatics Institute, Wellcome Genome Campus, Hinxton, Cambridge CB10 1SD, UK; Instituto de Biotecnología, Centro de Investigaciones en Ciencias Veterinarias y Agronómicas (CICVyA), Instituto Nacional de Tecnología Agropecuaria (INTA); Instituto de Agrobiotecnología y Biología Molecular (IABIMO), INTA-CONICET Nicolas Repetto y Los Reseros s/n (1686), Hurlingham, Buenos Aires, Argentina; Consejo Nacional de Investigaciones Científicas y Técnicas–CONICET, Ciudad Autónoma de Buenos Aires, Argentina

**Keywords:** Plant genomes, repetitive sequences, transposable elements, NLR genes, sequence masking, annotation

## Abstract

The annotation of repetitive sequences within plant genomes can help in the interpretation of observed phenotypes. Moreover, repeat masking is required for tasks such as whole-genome alignment, promoter analysis or pangenome exploration. While homology-based annotation methods are computationally expensive, k-mer strategies for masking are orders of magnitude faster. Here we benchmark a two-step approach, where repeats are first called by k-mer counting and then annotated by comparison to curated libraries. This hybrid protocol was tested on 20 plant genomes from Ensembl, using the kmer-based Repeat Detector (Red) and two repeat libraries (REdat and nrTEplants, curated for this work). We obtained repeated genome fractions that match those reported in the literature, but with shorter repeated elements than those produced with conventional annotators. Inspection of masked regions overlapping genes revealed no preference for specific protein domains. Half of Red masked sequences can be successfully classified with nrTEplants, with the complete protocol taking less than 2h on a desktop Linux box. The repeat library and the scripts to mask and annotate plant genomes can be obtained at https://github.com/Ensembl/plant-scripts.

## 1. Introduction

Besides genes, plant genomes contain intergenic sequences, with increasing repetitive sequences as genome size grows. The growth in repeat content is roughly linear up to a genome size of 10Gbp, including most known angiosperms, and then plateaus (1). The repetitive fraction of the genome is made up of low-copy repeats, simple repeats (such as satellite DNA), and transposable elements (TEs), which were discovered by Barbara McClintock in maize (2).

TEs can be important to explain observed phenotypes or domestication (see for instance reference (3)) and are used as a source of genetic variability in breeding programs (4). The hypothesis is that the copy-and-paste and cut-and-paste mechanisms of TEs might leave footprints in the genome and can potentially affect the expression, regulation or coding sequence of neighboring genes. Moreover, TEs are increasingly receiving attention in studies tackling plant pangenomes (see for instance (5)). According to the Wicker classification, plant TEs can be classified either as class I RNA retrotransposons or class II DNA transposons (6). Software resources such as RepeatMasker (RM) (7), RepBase (8) or RepetDB (9) that are typically used to annotate TEs in plant genomes use the Wicker classification rules (see (10) for a software review). However, it has been reported that disease resistance (R) genes, which are of great interest for plant breeding, are often masked by these annotation strategies (11). Furthermore, in our experience creating the Ensembl Plants resources, repeat annotation jobs can take up to several days on a computer cluster depending on the genome size. Moreover, the RepBase library requires subscription.

In addition to the intrinsic biological value of TEs, the annotation of repeats can be used to estimate assembly quality (12), as an alternative to gene completeness (13). For other genomic analyses, the bulk of repeated sequences may disrupt common computational genomic analyses and are thus often masked out without any classification attempt. For instance, whole-genome alignment (WGA), promoter analysis or the construction of graph genomes require the computation of resulting in frequency tables of k-mers, which are nucleotide words of size k. If repeated sequences are not masked, the frequency tables are severely biased and can affect the obtained results (see for instance (14)). While annotation approaches based on sequence similarity are computationally expensive, k-mer masking strategies are orders of magnitude faster (15–18) and, in our experience, much better to prepare WGAs of barley and wheat cultivars using LASTZ (19).

In this paper we benchmark a two-step approach for the annotation of plant repeated sequences. First, repeats are called by k-mer counting with the Repeat Detector (Red). Second, the discovered repeated sequences are annotated by sequence alignment to a newly curated metacollection of repeats called nrTEplants. We compare this approach to the conventional RM pipeline, with both nrTEplants and the REdat (20) library, on a set of 20 angiosperms from Ensembl. We then compare their performance and discuss the results. The nrTEplants library is bundled with scripts to mask and annotate genomes of angiosperms, enabling interoperability, reuse and reproducible analyses (21).

## 2. Materials and methods

### 2.1 Plant repeat libraries

We searched the literature for plant-specific libraries of repeated sequences and selected those in **Table 1**. While some are specific for a species or repeat family, others comprise repeats from mixed species, such as REdat from PGSB PlantsDB (20) or RepetDB (9). FASTA files with nucleotide sequences of repeats were downloaded from the indicated URLs or obtained from the authors.

**Table 1.**
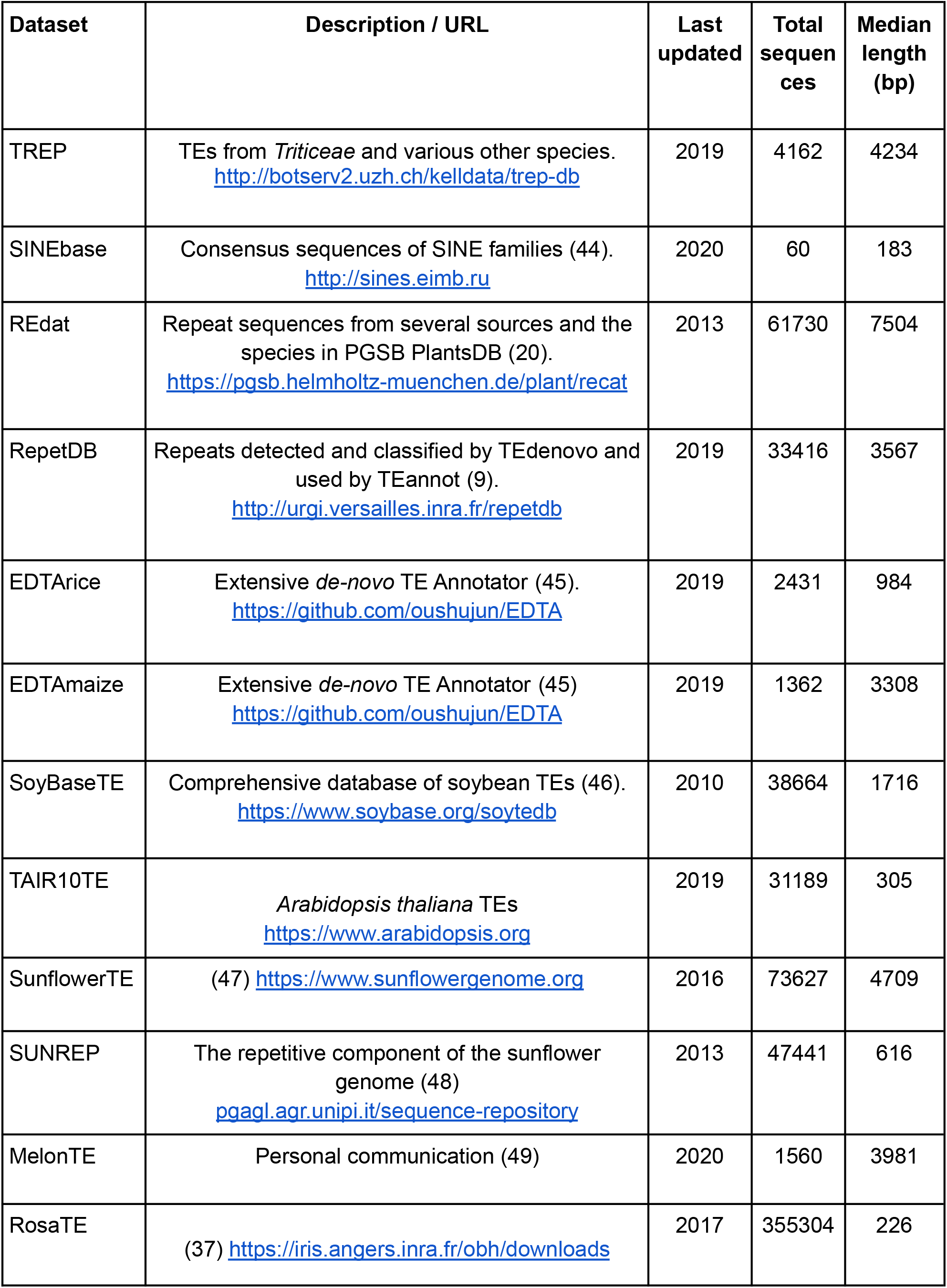
Collections of plant repeated sequences used as components of nrTEplants.

| Dataset | Description / URL | Last updated | Total sequences | Median length (bp) |
| --- | --- | --- | --- | --- |
| TREP | TEs from *Triticeae* and various other species.<br>http://botserv2.uzh.ch/kelldata/trep-db | 2019 | 4162 | 4234 |
| SINEbase | Consensus sequences of SINE families (44).<br>http://sines.eimb.ru | 2020 | 60 | 183 |
| REdat | Repeat sequences from several sources and the species in PGSB PlantsDB (20).<br>https://pgsb.helmholtz-muenchen.de/plant/recat | 2013 | 61730 | 7504 |
| RepetDB | Repeats detected and classified by TEdenovo and used by TEannot (9).<br>http://urgi.versailles.inra.fr/repetdb | 2019 | 33416 | 3567 |
| EDTArice | Extensive *de-novo* TE Annotator (45).<br>https://github.com/oushujun/EDTA | 2019 | 2431 | 984 |
| EDTAmaize | Extensive *de-novo* TE Annotator (45)<br>https://github.com/oushujun/EDTA | 2019 | 1362 | 3308 |
| SoyBaseTE | Comprehensive database of soybean TEs (46).<br>https://www.soybase.org/soytedb | 2010 | 38664 | 1716 |
| TAIR10TE | *Arabidopsis thaliana* TEs<br>https://www.arabidopsis.org | 2019 | 31189 | 305 |
| SunflowerTE | (47) https://www.sunflowergenome.org | 2016 | 73627 | 4709 |
| SUNREP | The repetitive component of the sunflower genome (48)<br>pgagl.agr.unipi.it/sequence-repository | 2013 | 47441 | 616 |
| MelonTE | Personal communication (49) | 2020 | 1560 | 3981 |
| RosaTE | (37) https://iris.angers.inra.fr/obh/downloads | 2017 | 355304 | 226 |

### 2.2 Plant cDNA sequences

Plant species in Ensembl Plants release 46 (November 2020) (22) were ranked in terms of the number of proteins reviewed in Uniprot by February 22nd, 2020 (23). This was considered as an indicator of annotation quality. A list of the best annotated dicot and monocot species was produced, including *Arabidopsis thaliana, Brassica napus, Glycine max, Helianthus annuus, Medicago truncatula, Phaseolus vulgaris, Populus trichocarpa, Solanum lycopersicum, Vitis vinifera, Brachypodium distachyon, Hordeum vulgare, Oryza sativa* subsp. *japonica, Sorghum bicolor* and *Zea mays*. Transcripts from these species were downloaded with the script *ens_sequences.pl* from https://github.com/Ensembl/plant-scripts.

### 2.3 Sequence clustering

All cDNA and TE sequences were clustered with GET_HOMOLOGUES-EST version 10042020 (24). This software runs BLASTN and the MCL algorithm, and computes coverage by combining local alignments. The sequence identity cutoff was 95% and the alignment coverage 75%. Global variables in script *get_homologues-est.pl* lines L36-7 were set to $MAXSEQLENGTH = 55000 and $MINSEQLENGTH = 90. Sequences were clustered with command *get_homologues-est.pl -d repeats -m cluster -M -t 0 -i 100*. Check https://github.com/Ensembl/plant_tools/tree/master/bench/repeat_libs for more details.

### 2.4 Positive control Pfam domains

A list of 22 Pfam domains found in transposable elements was curated (25), available at https://github.com/Ensembl/plant_tools/blob/master/bench/repeat_libs/control_pos.list

### 2.5 Negative control: Pfam domains of disease resistance (R) genes

For the identification and curation of Pfam domains encoded by R genes, the following steps were performed. First, a set of 153 protein sequences encoded by reference R genes (i.e. cloned and/or with robust evidence) was retrieved from http://www.prgdb.org/prgdb (26). Second, the program *hmmscan* from HMMER v3.2.1 (27) was used for initial Pfam domain identification (v32, *default* settings), yielding a total of 60 Pfam hidden Markov models. The observed order and combinations of Pfam domains were retrieved. Third, the proteins of 6 plant species (*A. thaliana, B. distachyon, G. max, H. annuus, H. vulgare* and *Triticum aestivum*) containing at least one of the 60 pfam previously identified were retrieved from https://plants.ensembl.org/biomart/martview (28). These proteins were subsequently filtered, retaining only those with the ordered combinations of Pfam domains observed in the reference R proteins, and were considered as potential R proteins (428 in *A. thaliana*, 577 in *B. distachyon*, 1,008 in *G. max*, 849 in *H. annuus*, 838 in *H. vulgare*, and 3,607 in *T. aestivum*). From the initial set of Pfam domains, only 43 were consistently identified in our final panel of potential R gene’s encoded proteins, and used as negative control. Note that one of them (PF02892, zf-BED) is often found in transposases (25). The list is available at https://github.com/Ensembl/plant_tools/blob/master/bench/repeat_libs/control_neg_NLR.list.

### 2.6 De novo annotation of nucleotide-binding and leucine-rich repeat immune receptor (NLR) genes

The NLR-annotator software was used for *de novo* annotation of NLR genes, which are the most abundant R genes characterized to date, in whole genome sequences (29). Briefly, the set of 20 plant genomes were dissected into fragments of 20 kb length, with 5 kb overlaps, using the *ChopSequence.jar* routine. The cut sequences were then scanned to find NLR-associated sequence motifs using *NLR-Parser.jar*. Finally, *NLR-Annotator.jar* was used to integrate the annotated motifs and retrieve the actual NLR loci in BED format. In order to compute intersections with repeats only NLR loci with overlap > 50bp were considered. Moreover, to account for the fact that the tested masking strategies cover different fractions of the genome, odd ratios of NLR masking were computed with **Equation 1**:

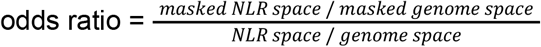

### 2.7 Masking and annotation of repeats in plant genomes

RepeatMasker version v4.0.5 and a fork of Repeat Detector (Red) v2.0 adapted for Ensembl, available at https://github.com/EnsemblGenomes/Red, were used to call repeats in plant genomes with libraries REdat v9.3 and nrTEplanst v0.3. Red was called from script https://github.com/Ensembl/plant-scripts/blob/master/repeats/Red2Ensembl.py, which can run several sequences in parallel and feed the results into a Ensembl core database (30). In addition, minimap2 version 2.17-r974-dirty (31) was used to annotate repeats called by Red with sequences from nrTEplants as follows: *minimap2 -K100M --score-N 0 -x map-ont nrTEplants*. Minimap2 is called from script https://github.com/Ensembl/plant-scripts/blob/master/repeats/AnnotRedRepeats.py, which parses its output to annotate the repeats. By default only repeats with length > 90 bp are processed. TE classification terms are parsed from the FASTA header of the library after a hash char (#, i.e. ‘RLG_43695:mipsREdat_9.3p_ALL#LTR/Gypsy’). Elapsed runtime and RAM consumption was obtained with *command time -v*.

Genomic intersections among repeated sequences called by Red and RM and genomic features (ie protein-coding genes, exons, proximal downstream/upstream 500bp windows, NLR loci) were computed with Bedtools (v2.26.0) (32) using *bedtools intersect -a bed/genes.bed -b repeat.bed -sorted -wo*. To avoid redundancy, exons were extracted from Ensembl canonical transcripts (see http://plants.ensembl.org/info/website/glossary.html). Neighbor genes were also subtracted from downstream/upstream windows.

### 2.8 K-mer analysis of repeats in downstream/upstream windows

Repeats overlapping proximal downstream/upstream 500bp windows were extracted using *bedtools intersect* analysis and the sequences cut with *bedtools getfasta*. Canonical K-mers with K=[16,21,31] were counted with Jellyfish v2.3.0 (33) with commands *jellyfish-linux count -C -m K -s 2G -t 4* and *jellyfish-linux dump -L 20*.

### 2.9 Enrichment of Pfam domains

Enrichment was computed with R function fisher.test (34) and Pfam domains (25) retrieved with recipe B4 of https://github.com/Ensembl/plant-scripts (35). Pfam domain counts for the complete proteome were used as expected frequencies. Only genes with an overlap > 50bp and domains with False Discovery Rate adjust values (p<0.05) were considered.

### 2.10 Control sets of annotated repeated sequences

Repeated sequences annotated by sequencing consortia of olive tree (36), *Rosa chinensis* (37) and sunflower (38) were downloaded and formatted from https://genomaolivar.dipujaen.es/db/downloads.php, https://iris.angers.inra.fr/obh/downloads and https://sunflowergenome.org/annotations-data.

## 3. Results and discussion

### 3.1 Construction and benchmark of a non-redundant library of repeats: nrTEplants

A set of plant TE libraries and annotated repeats from selected species, listed on **Table 1**, plus cDNA sequence sets from the best annotated plant species in Ensembl, were curated and their TE classification terms uniformized. Then, they were merged and clustered (95% identity, 75% coverage of shortest sequence). From the resulting 994,349 clusters, a total 174,426 clusters contained TE sequences and were 6-frame translated and assigned Pfam domains. Of these, a subset of 8,910 mixed clusters comprised both TE and cDNA sequences and required further processing (see example on **Figure S1**). After empirical assessment, we decided to take only clusters i) containing sequences from at least 6 different TE libraries (6 replicates), which eventually left out RosaTE repeats; and ii) with a fraction of sequences marked as ‘Potential Host Gene’ in RepetDB < 0.00. The resulting nrTElibrary contains 171,104 sequences (see **Tables S1 and S3**).

In order to benchmark the newly constructed library we compiled a positive control, comprising 22 Pfam domains found in transposable elements, and a negative control, a list of 43 Pfam domains found in disease resistance NLR genes. With these controls, we estimated the sensitivity (0.909) and specificity (0.947). The nrTEplants library can be obtained at https://github.com/Ensembl/plant-scripts/releases/tag/v0.31.

### 3.2 Masking repeats within plant genomes

A set of 20 plant genomes were selected from Ensembl (22) to benchmark repeat calling strategies. These are listed on **Table 2** next to the genomic fraction of repeats reported in the literature and their GC content. All these genome sequences were annotated with RepeatMasker (7), nrTEplants and REdat (20), a repeat library used in Ensembl Plants. In addition, the fraction of repeats called by Red, based on K-mer enrichment, is also shown. Note that Red automatically selected k values from 13 to 16.

**Table 2.** Plant genomes from release 49 (September 2020) of Ensembl Plants (22) used in this work and their reported repeated fractions in the literature.

| Species | %GC | Assembled genome size (Mbp) | Reported % repeated fraction (source) |
| --- | --- | --- | --- |
| *Arabidopsis thaliana* | 36.1 | 119.7 | 19.0 (50) |
| *Arabidopsis halleri* | 36.0 | 196.2 | 32.7 (50) |
| *Prunus dulcis* | 37.6 | 227.5 | 37.6 (51) |
| *Brachypodium distachyon* | 46.4 | 271.2 | 21.4 (52) |
| *Brassica rapa* | 35.3 | 283.8 | 32.3 (53) |
| *Trifolium pratense* | 32.4 | 304.8 | 41.8 (54) |
| *Arabis alpina* | 36.8 | 308.0 | 47.9 (55) |
| *Cucumis melo* | 33.5 | 357.9 | 44.0 (56) |
| *Citrullus lanatus* | 33.6 | 365.5 | 45.2 (57) |
| *Oryza sativa* | 43.6 | 375.0 | 35 (58) |
| *Setaria viridis* | 46.2 | 395.7 | 46 (59) |
| *Vitis vinifera* | 34.5 | 486.3 | 41.4 (60) |
| *Rosa chinensis* | 38.8 | 515.6 | 67.9 (61) |
| *Camelina sativa* | 36.6 | 641.4 | 28 (62) |
| *Malus domestica* | 38.0 | 702.9 | 59.5 (63) |
| *Olea europaea* | 35.4 | 1140.9 | 43 (64) |
| *Zea mays* | 46.9 | 2135.1 | 85 (65) |
| *Helianthus annuus* | 38.5 | 3027.8 | 74.7 (38) |
| *Aegilops tauschii* | 46.3 | 4224.9 | 85.9 (66) |
| *Triticum turgidum* | 46.0 | 10463.1 | 82.2 (67) |

On **Figure 1** the resulting percentages of repeated sequences are plotted next to the values reported in the literature. The median difference between REdat repeated fraction and the literature reports is 26.5%. This number is 9.8% for nrTEplants and 4.3% for Red. These results suggest that Red can successfully mask any genomes without previous knowledge of the repetitive sequence repertoire of a species. Moreover, repeats called by Red generally overlap sequences masked with REdat (66.6%) and nrTEplants (73.8%).

**Figure 1.**
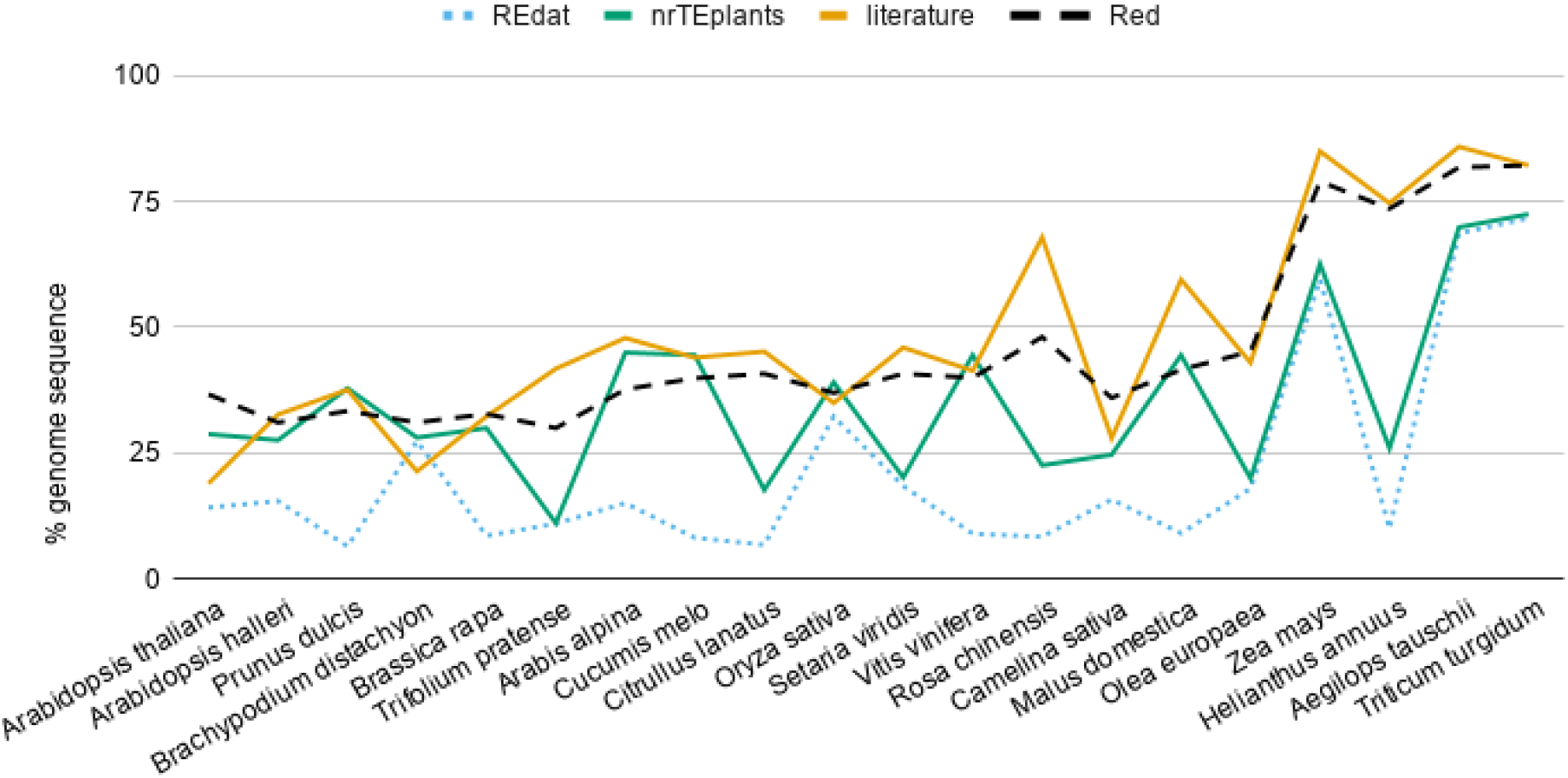
Fraction of repeated sequences in plant genomes. Twenty genomes from release 49 (November 2020) of Ensembl Plants were annotated with RepeatMasker (7) and libraries REdat (20) and nrTEplants. The resulting percentage of repeated sequences is plotted next to the values reported in the literature for those genomes and the fraction of repeats provided by Repeat Detector (Red), based on K-mer enrichment (15). Species are sorted from small to large genome size.

**Table 3** summarizes the number of repeats, and their length, called by all tested strategies. We observe that Red calls more (a median of 845 per Mbp, compared to 391 for nrTEplants and 221 for REdat) but shorter repeats (median 143, compared to 284 and 217, respectively). Note that the numbers below are shown in the same REd, nrTEplants, REdat order for clarity.

**Table 3.**
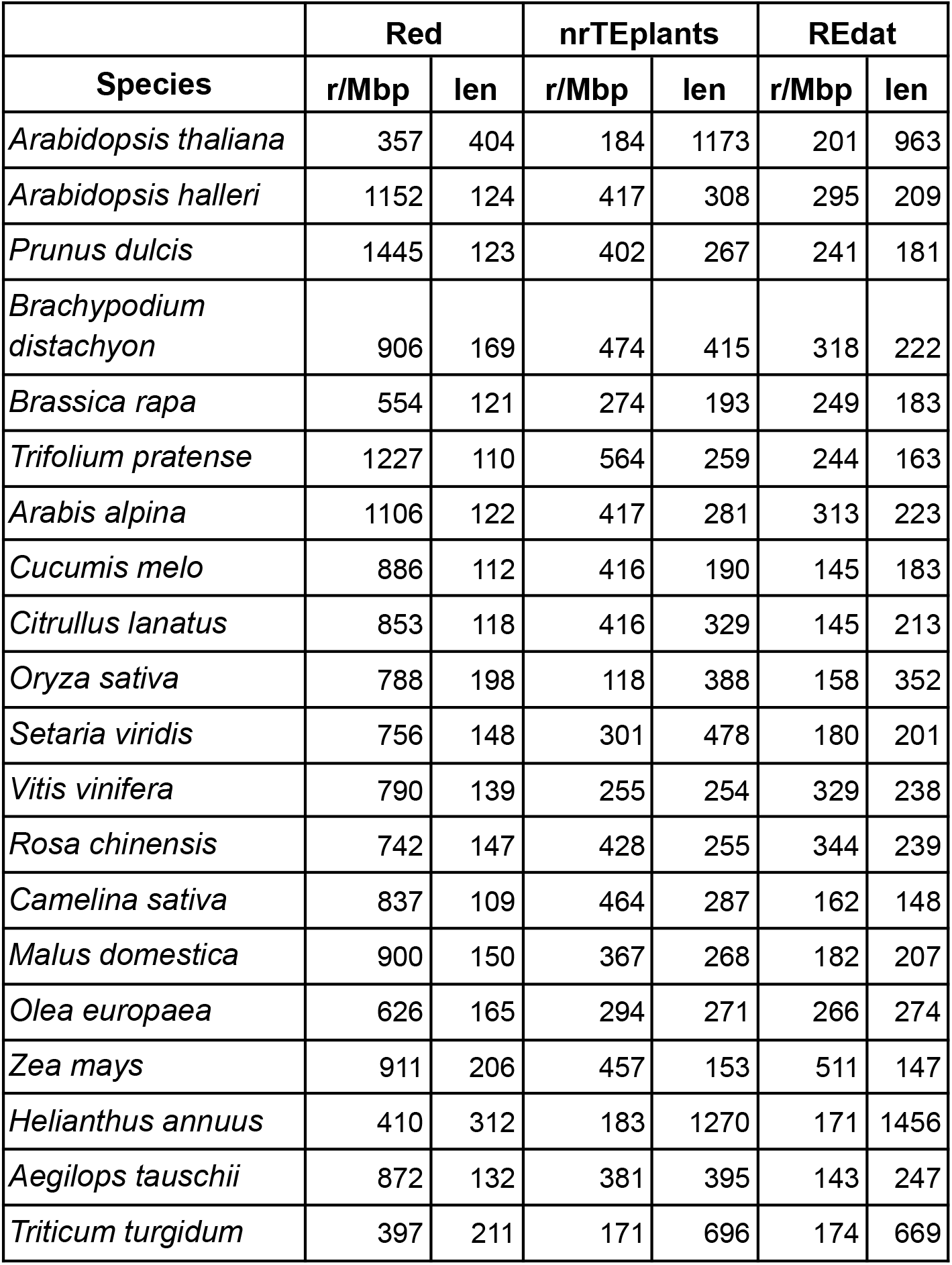
Summary of repeated sequences annotated with Red (15) and RepeatMasker (7) with libraries nrTEplants and REdat (20). Repeats per Mbp (r/Mbp) and median length (len) are shown.

|  | Red |  | nrTEplants |  | REdat |  |
| --- | --- | --- | --- | --- | --- | --- |
| Species | r/Mbp | len | r/Mbp | len | r/Mbp | len |
| *Arabidopsis thaliana* | 357 | 404 | 184 | 1173 | 201 | 963 |
| *Arabidopsis halleri* | 1152 | 124 | 417 | 308 | 295 | 209 |
| *Prunus dulcis* | 1445 | 123 | 402 | 267 | 241 | 181 |
| *Brachypodium distachyon* | 906 | 169 | 474 | 415 | 318 | 222 |
| *Brassica rapa* | 554 | 121 | 274 | 193 | 249 | 183 |
| *Trifolium pratense* | 1227 | 110 | 564 | 259 | 244 | 163 |
| *Arabis alpina* | 1106 | 122 | 417 | 281 | 313 | 223 |
| *Cucumis melo* | 886 | 112 | 416 | 190 | 145 | 183 |
| *Citrullus lanatus* | 853 | 118 | 416 | 329 | 145 | 213 |
| *Oryza sativa* | 788 | 198 | 118 | 388 | 158 | 352 |
| *Setaria viridis* | 756 | 148 | 301 | 478 | 180 | 201 |
| *Vitis vinifera* | 790 | 139 | 255 | 254 | 329 | 238 |
| *Rosa chinensis* | 742 | 147 | 428 | 255 | 344 | 239 |
| *Camelina sativa* | 837 | 109 | 464 | 287 | 162 | 148 |
| *Malus domestica* | 900 | 150 | 367 | 268 | 182 | 207 |
| *Olea europaea* | 626 | 165 | 294 | 271 | 266 | 274 |
| *Zea mays* | 911 | 206 | 457 | 153 | 511 | 147 |
| *Helianthus annuus* | 410 | 312 | 183 | 1270 | 171 | 1456 |
| *Aegilops tauschii* | 872 | 132 | 381 | 395 | 143 | 247 |
| *Triticum turgidum* | 397 | 211 | 171 | 696 | 174 | 669 |

**Figure 2** summarizes how called repeats overlap with genes, exons and 500pb windows upstream and downstream genes. It can be seen that Red overlaps more with genes (median 17.8%, compared to 13.7% and 5%, respectively), but when exons are considered these numbers change to 9%, 15% and 4%, respectively. The figure also shows that Red masks more proximal upstream and downstream space, which will likely have a positive impact on k-mer counting strategies for promoter analysis (39). The analysis on **Table S4** shows that Red identifies four times more K-mers with 20+ copies in this regulatory space, which agrees with recent work that found that unidentified TEs are over-represented in specific regulatory networks (40).

**Figure 2.**
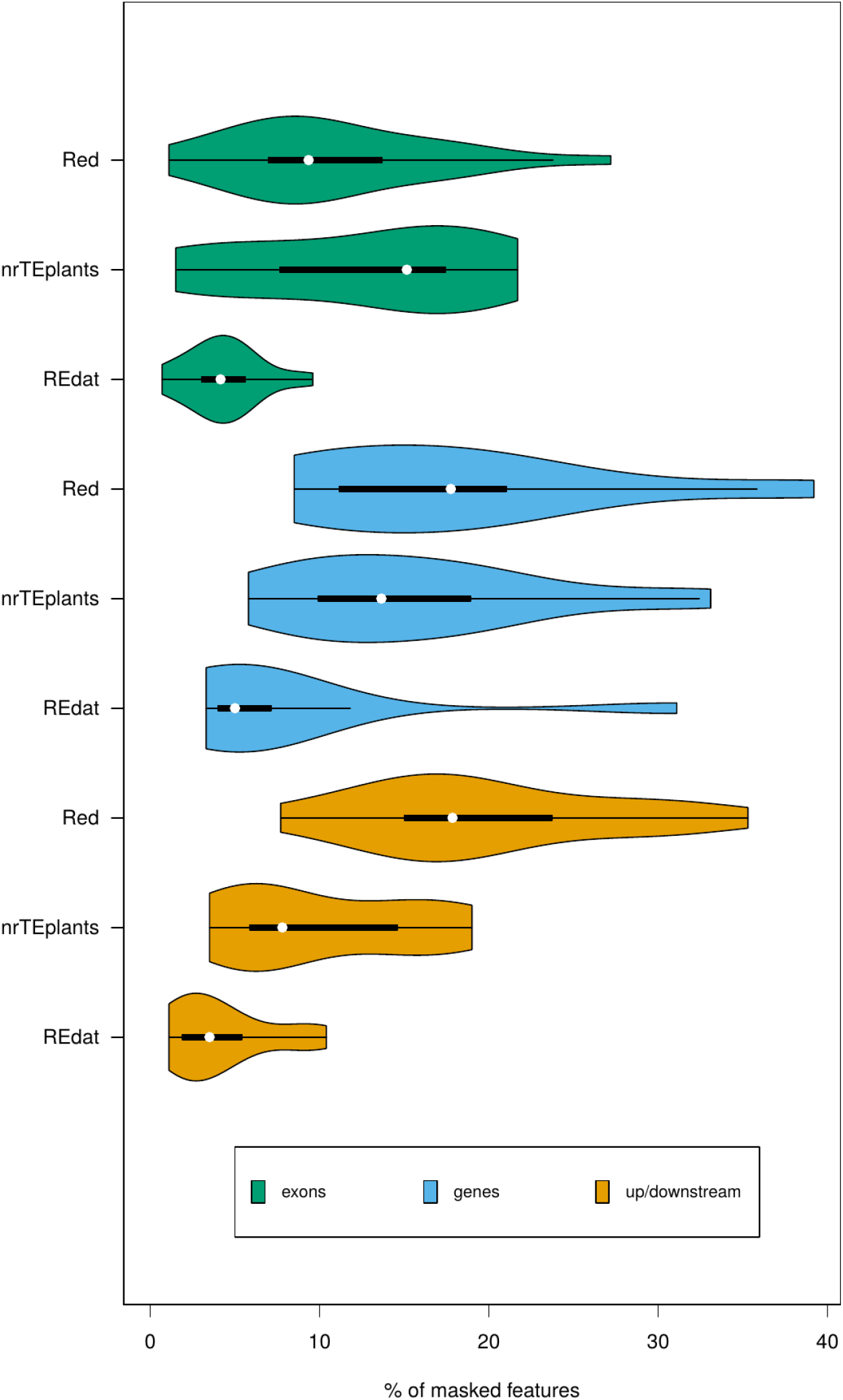
Fraction of exons, genes and 500bp up/downstream regions overlapping annotated repeats in plant genomes. Twenty genomes from release 49 (November 2020) of Ensembl Plants were annotated with Red (15) or RepeatMasker (7) with libraries REdat (20) and nrTEplants. The median genome space occupied by genes, exons and proximal upstream/downstream 500bp windows in these species are 126.7Mbp, 62.9Mbp and 37.7Mbp, respectively. This plot was generated with R package vioplot (42).

In order to check whether the compared approaches masked preferentially genes from certain families, a Pfam enrichment analysis was carried out and summarized on **Figure 3**. It can be seen that Red repeats show the least enrichment. Furthermore, Red enriched Pfam domains are different across genomes with the exception of four domains found enriched in three genomes (reverse transcriptase-like, TIR, NB-ARC, and integrase core domains). In contrast, a few Pfam domains were enriched in 10+ genomes in genes overlapping repeats annotated with RM. The results on **Table S5** show that nrTEplants performs better than REdat in this respect, with 87 domains compared to 153 (Red had 39 in total).

**Figure 3.**
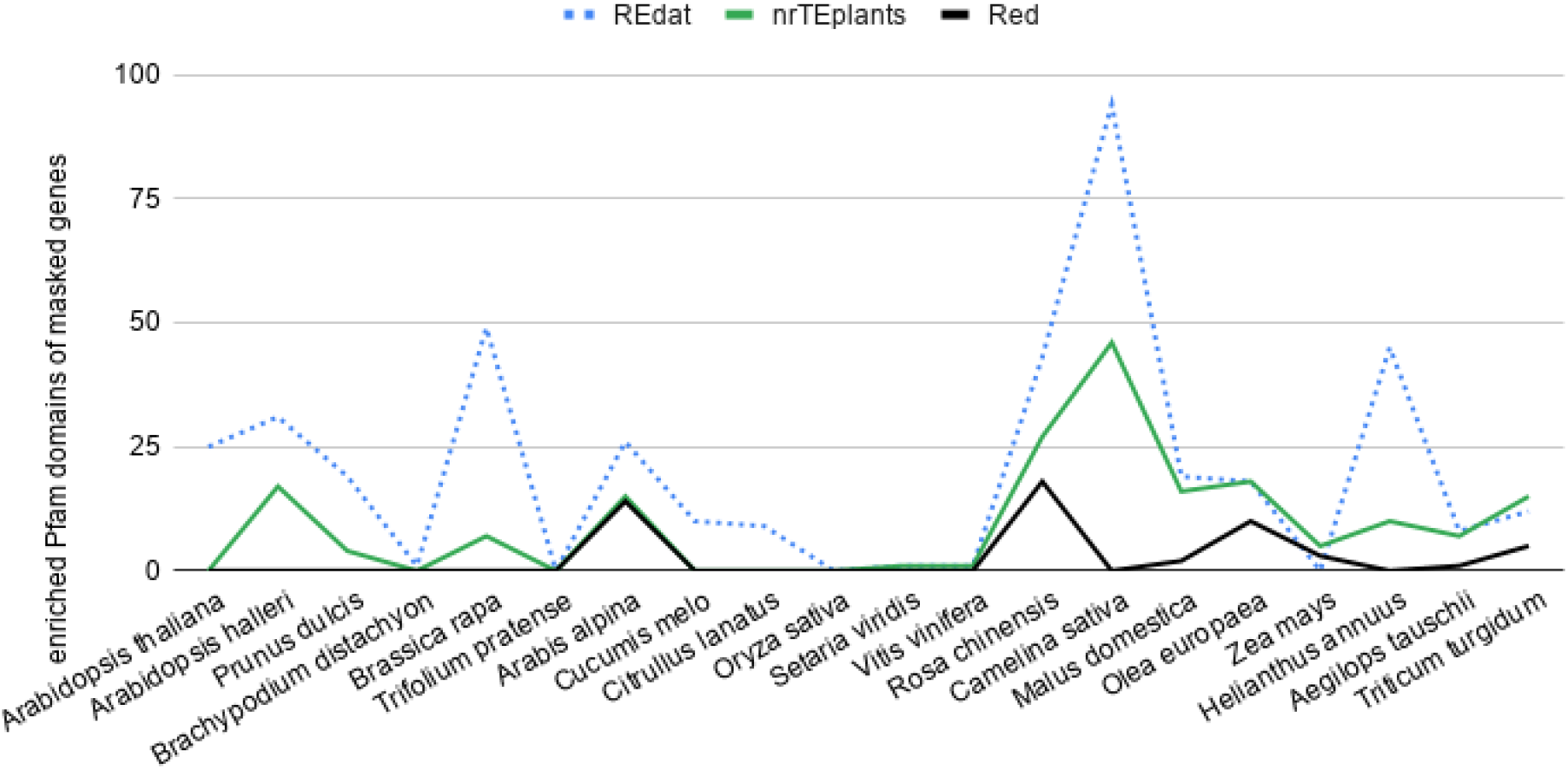
Enriched Pfam domains of protein-coding genes overlapping repeats. Twenty genomes from release 49 (November 2020) of Ensembl Plants were annotated with Repeat Detector (Red) (15) and RepeatMasker (7) with libraries REdat (20) and nrTEplants.

As gene annotation is frequently performed after repeat masking, we reasoned this could affect the Pfam enrichment analyses. Therefore we carried out a complementary analysis where NLR genes were called *de novo* on the genomic sequences instead of using the Ensembl gene annotation. The results, summarized on **Table S5**, confirm that Red tends to mask less NLR genes than expected at genomic scale, with only one species (*Trifolium pratense*) with an odds ratio > 1. In contrast, we obtained odd ratios greater than 1 for several species with REdat (n=7) and nrTEplants (n=12).

Overall, we observe that Red consistently produces repeated fractions similar to the expected values from the literature. The library nrTEplants also shows good performance in most species, but fails to recover the expected repeat fraction in cases such as melon or sunflower. Red also performs better than nrTEplants in terms of exon and upstream/downstream overlap. Furthermore, Red does not seem to systematically mask certain gene families. The lower values observed for REdat are a consequence of it underpredicting repeats, showing less sensitivity than the other approaches.

### 3.3 Annotating Red-masked repeats within genomes with nrTEplants and minimap2

In the previous analyses we showed that Red-base masking is an effective way of calling repeats in plant genomes. Moreover, we observed that nrTEplants behaves better than REdat in most cases. Therefore, we wanted to check whether repeats called with Red can be annotated and classified. For that, we aligned the repeat sequences against the non-redundant nrTElibrary with minimap2. The results are plotted on **Figure 4**, where it can be seen that, in most species, more than half of the repeat space can be annotated (median 65.9%). As our library contains only transposable elements, we expected a fraction of unmapped space containing simple repeats or satellite DNA. However, as also observed on **Figure 1**, in a few species, only a small repeat fraction could be classified. We reasoned this was due to repeat consensus not represented in the library. This was confirmed in a separate experiment where olive and *R. chinensis* repeated sequences obtained from their authors were mapped to Red repeats, as seen in **Figure 4** in a grey dashed line. A positive control was also carried out with sunflower repeated sequences, in order to confirm that no valuable repeats had been lost during the construction of nrTEplants.

**Figure 4.**
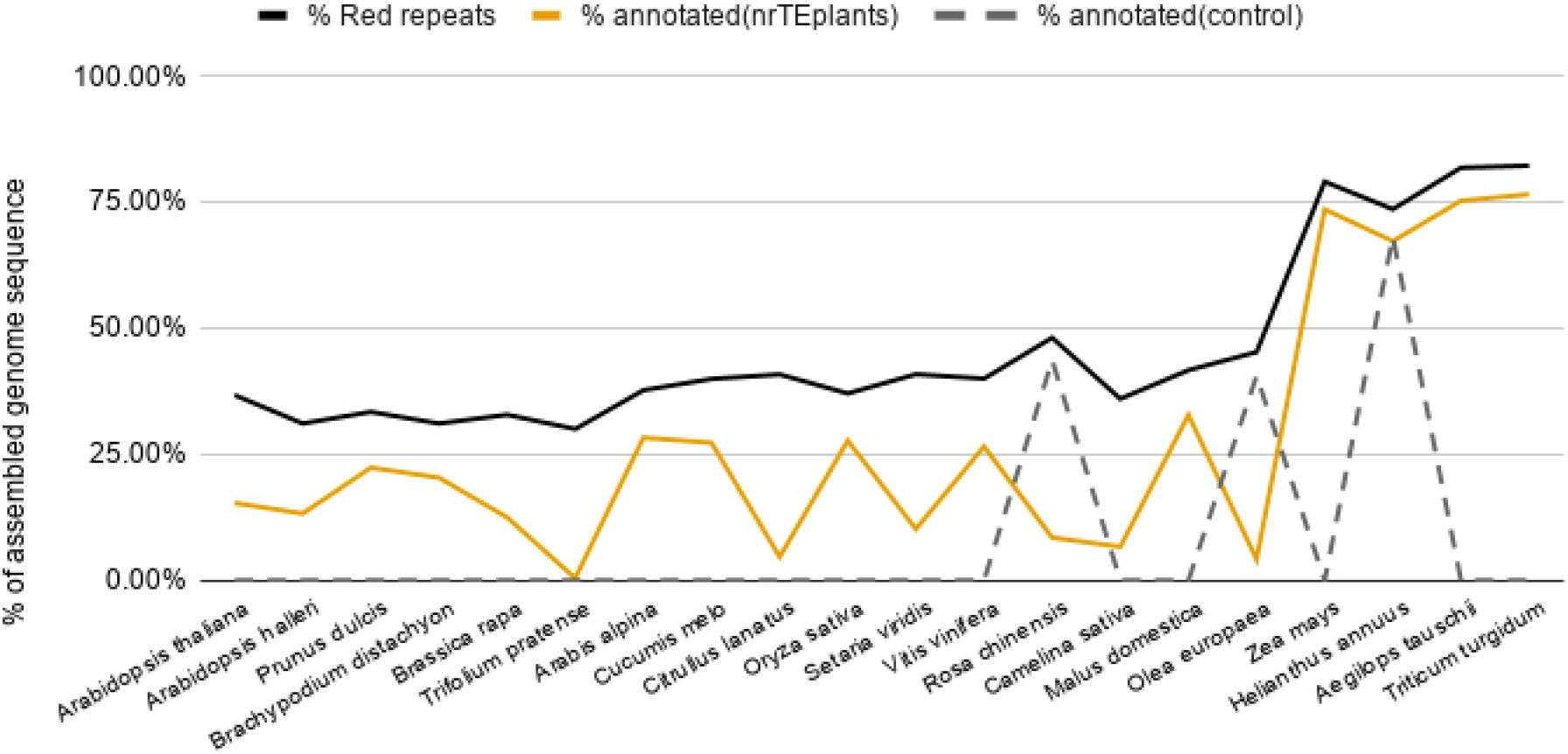
Fraction of Red repeats mapped to nrTEplants sequences. Twenty genomes from release 49 (November 2020) of Ensembl Plants were annotated with Repeat Detector (15). The resulting repeats were subsequently mapped to library nrTEplants with minimap2 (31), producing the genome fractions shown. Repeats from three species (*R. chinensis, O. europaea* and *H. annuus*) were also mapped to annotated repeats provided by the respective sequencing consortia as a control. Species are sorted from small to large genome size.

These results were obtained with the default *map-ont* settings of minimap2. We also tried *map-pab* and *asm20* settings, but obtained similar results. Red clover (*Trifolium pratense*) was re-analyzed replacing minimap2 with BLAST algorithms *megablast, dc-megablast, blastn* and *rmblastn* (41). Compared to the mapped fraction produced by minimap2 (0.4%), a maximum value of 6.1% was obtained with blastn. This modest gain in sensitivity required 1,412 minutes. The algorithm *rmblastn*, used by RM, yielded a mapped fraction of 0.7%. We concluded that the alternatives to minimap2 offered little gain at the cost of spiralling computing time.

**Figure 5** shows the runtime and RAM required by the two-step protocol presented in this paper, measured on a CentOS7.9 computer using 4 cores of a Xeon E5-2620 v4 (2.10GHz) CPU. Panels A and B correspond to the first step, Red masking. It can be seen that all genomes tested take less than 40 min to run, with the exception of tetraploid *Triticum turgidum*, which took 71 min. The memory consumption was below 20GB in all cases, but climbed to 22.7GB and 29.9GB in *A. tauschii* and *T. turgidum*. Panel C illustrates the runtime of the second step, the mapping of nrTEplants. It can be seen that all plants required less than 27 min, except *A. tauschii* and *T. turgidum*, that took 3h and 1h respectively. The memory consumed by minimap2 was 3.8-4.0GB in all cases.

**Figure 5.**
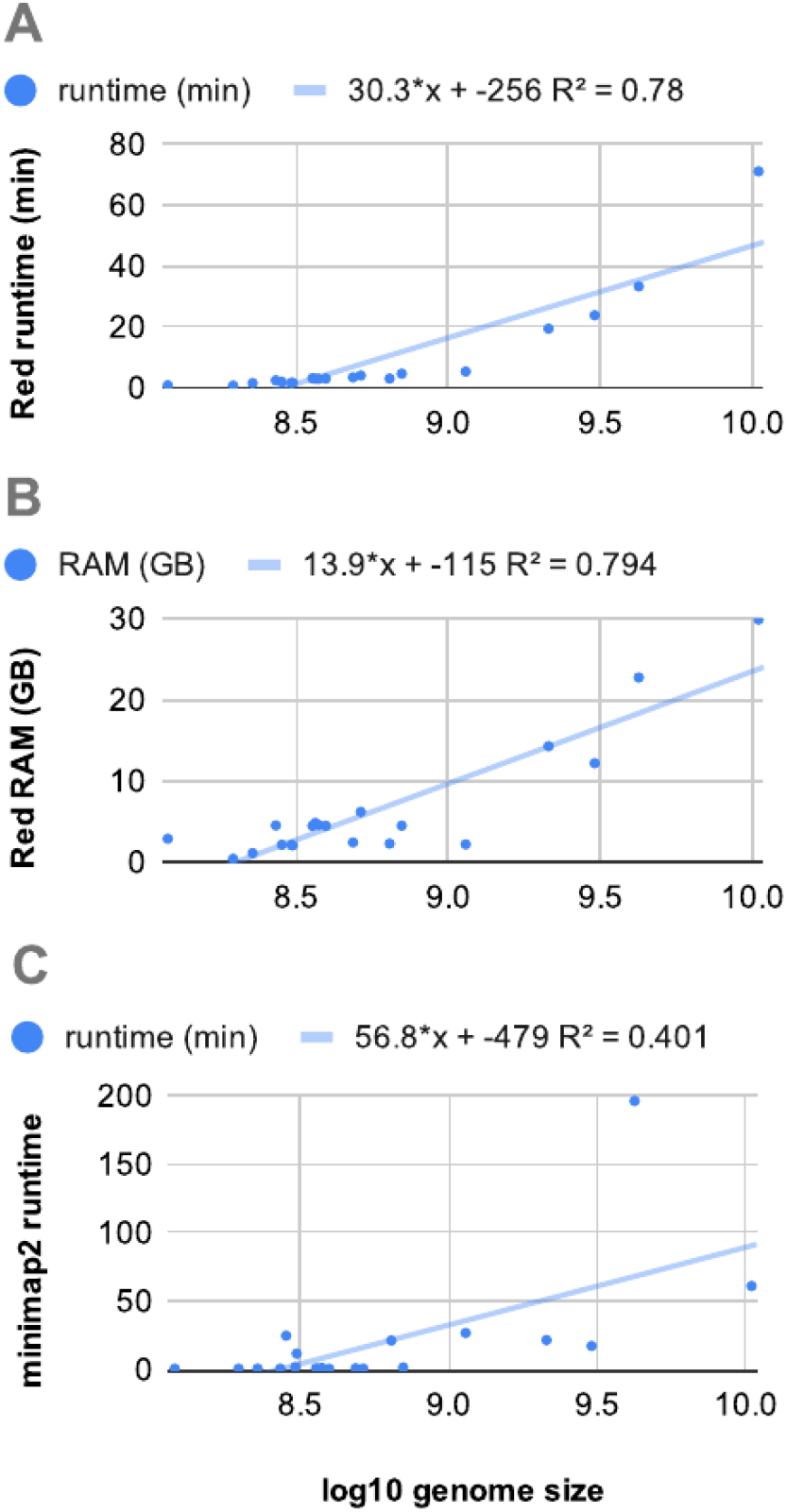
Runtime and memory requirements of a two-step repeat annotation protocol based on the Repeat Detector (15), minimap2 (31) and the nrTEplants library. The protocol was tested on twenty genomes from release 49 (November 2020) of Ensembl Plants. Similar values were measured on a Ubuntu box with four Core i5-6600 (3.30GHz) CPU cores.

## Conclusions

The hybrid, two-step methodology presented in this paper was tested on 20 angiosperms with genome sizes from 0.12 to 10.46 Gbp. Compared to RM, Red calls more and shorter repeats, and the obtained repeated genome fractions agree closely with those reported in the literature. Moreover, we find that Red k-mer masking does not have a preference for particular protein-coding families, in contrast to repeats annotated with RM using REdat and nrTEplants. Overall, more than half of Red masked sequences can be classified with nrTEplants, except in species with repeats not present in that library. Our protocol takes less than 2h to run and up to 30GB of RAM, and can use nrTEplants or any repeat library in FASTA format.

## Acknowledgements

We are grateful to Hani Girgis for his help with the Red source code and Doreen Ware for comments on drafts of this manuscript. We thank the Gramene team for continuous support and cooperation, as well as members of the Ensembl team for developing and maintaining the front-end and back-end software and infrastructure that underpins Ensembl Plants.

## Funding

The UK Biosciences and Biotechnology Research Council [BB/P016855/1, BB/P027849/1 and Ensembl-4-Breeders workshop support], the National Sciences Foundation [1127112], the ELIXIR implementation studies FONDUE and ‘Apple as a Model for Genomic Information Exchange’ and the European Molecular Biology Laboratory. Funding for open access charge: UK Biosciences and Biotechnology Research Council [BB/P016855/1]. Conflict of interest statement. Paul Flicek is a member of the Scientific Advisory Boards of Fabric Genomics, Inc. and Eagle Genomics, Ltd.

## Supplementary Figures

**Figure S1.**
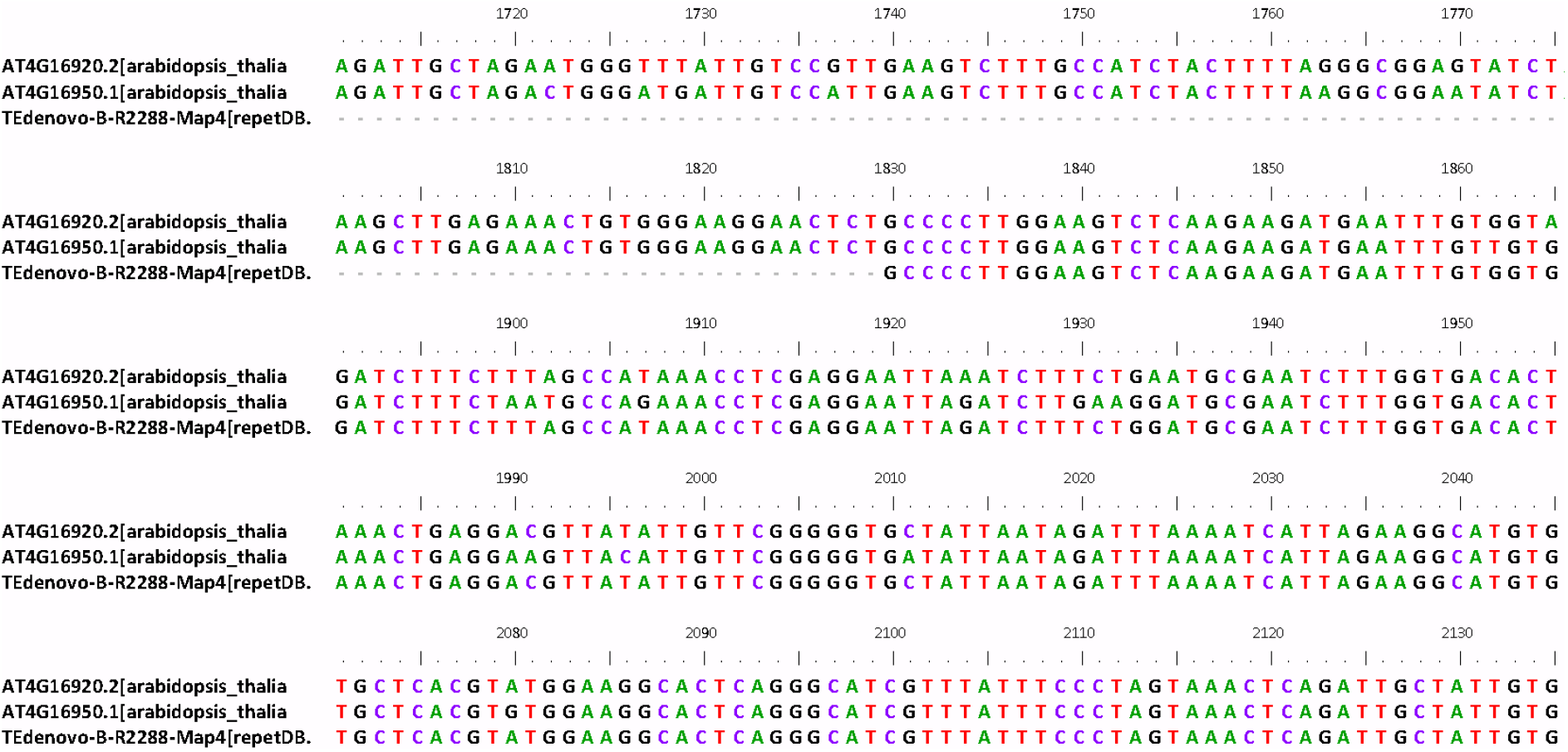
Cluster with two *Arabidopsis thaliana* cDNA sequences (AT4G16920.2 and AT4G16950.1) and transposable element TEdenovo-B-R2288-Map4 from library repetDB. These sequences contain Pfam domain NB-ARC (PF00931), which is part of NLR defense proteins. Figure generated with Bioedit (43) from cluster 269_AT4G16920.2.

## Supplementary Tables

**Table S1.** Contribution of individual datasets to the nrTEplants library of repeats (v0.3).

| Dataset | # of sequences |
| --- | --- |
| SunflowerTE | 58071 |
| REdat | 39509 |
| repetDB | 26006 |
| SoyBaseTE | 21766 |
| TAIR10TE | 21056 |
| EDTArice | 1844 |
| MelonTE | 1171 |
| EDTAmaize | 805 |
| TREP | 789 |
| SINEbase | 44 |
| SUNREP | 43 |

**Table S2.** Enriched Pfam domains of protein-coding genes overlapping repeats called with Red (15) and RepeatMasker (7) with libraries nrTEplants and REdat (20). Only domains found enriched in at least three species are shown. Domains in bold are shared by all repeat-calling strategies and correspond to Integrase core domains (PF00665), NB-ARC (PF00931), Reverse transcriptase-like (PF13456) and TIR (PF01582).

| REdat |  | nrTEplants |  | Red |  |
| --- | --- | --- | --- | --- | --- |
| Pfam domain | count | Pfam domain | count | Pfam domain | count |
| PF00069 | 13 | **PF00931** | 11 | **PF13456** | 3 |
| PF07714 | 10 | PF07727 | 6 | **PF01582** | 3 |
| PF00005 | 10 | PF07714 | 6 | **PF00931** | 3 |
| PF01095 | 9 | **PF01582** | 6 | **PF00665** | 3 |
| PF13947 | 8 | PF00078 | 6 |  |  |
| PF07727 | 8 | PF00069 | 6 |  |  |
| **PF00931** | 8 | PF18052 | 5 |  |  |
| PF13855 | 7 | PF14223 | 5 |  |  |
| PF01453 | 7 | PF07725 | 5 |  |  |
| PF00847 | 7 | **PF00665** | 5 |  |  |
| **PF00665** | 7 | PF17921 | 4 |  |  |
| PF00078 | 7 | PF13976 | 4 |  |  |
| PF14510 | 6 | PF13855 | 4 |  |  |
| PF14432 | 6 | PF03732 | 4 |  |  |
| PF14223 | 6 | PF00560 | 4 |  |  |
| PF08370 | 6 | PF14111 | 3 |  |  |
| PF08276 | 6 | PF13947 | 3 |  |  |
| **PF01582** | 6 | **PF13456** | 3 |  |  |
| PF00954 | 6 | PF08276 | 3 |  |  |
| PF00067 | 6 | PF08263 | 3 |  |  |
| PF17921 | 5 | PF01453 | 3 |  |  |
| PF17919 | 5 | PF00954 | 3 |  |  |
| PF17917 | 5 | PF00122 | 3 |  |  |
| PF13976 | 5 | PF00005 | 3 |  |  |
| PF11721 | 5 |  |  |  |  |
| PF08387 | 5 |  |  |  |  |
| PF08284 | 5 |  |  |  |  |
| PF08263 | 5 |  |  |  |  |
| PF07725 | 5 |  |  |  |  |
| PF07723 | 5 |  |  |  |  |
| PF03732 | 5 |  |  |  |  |
| PF00560 | 5 |  |  |  |  |
| PF00082 | 5 |  |  |  |  |
| PF19055 | 4 |  |  |  |  |
| PF17766 | 4 |  |  |  |  |
| PF08246 | 4 |  |  |  |  |
| PF02365 | 4 |  |  |  |  |
| PF01061 | 4 |  |  |  |  |
| PF00664 | 4 |  |  |  |  |
| PF00201 | 4 |  |  |  |  |
| PF00112 | 4 |  |  |  |  |
| PF00098 | 4 |  |  |  |  |
| PF00012 | 4 |  |  |  |  |
| PF14380 | 3 |  |  |  |  |
| **PF13456** | 3 |  |  |  |  |
| PF12819 | 3 |  |  |  |  |
| PF08372 | 3 |  |  |  |  |
| PF08031 | 3 |  |  |  |  |
| PF05922 | 3 |  |  |  |  |
| PF00450 | 3 |  |  |  |  |
| PF00305 | 3 |  |  |  |  |
| PF00249 | 3 |  |  |  |  |
| PF00223 | 3 |  |  |  |  |

**Table S3.**
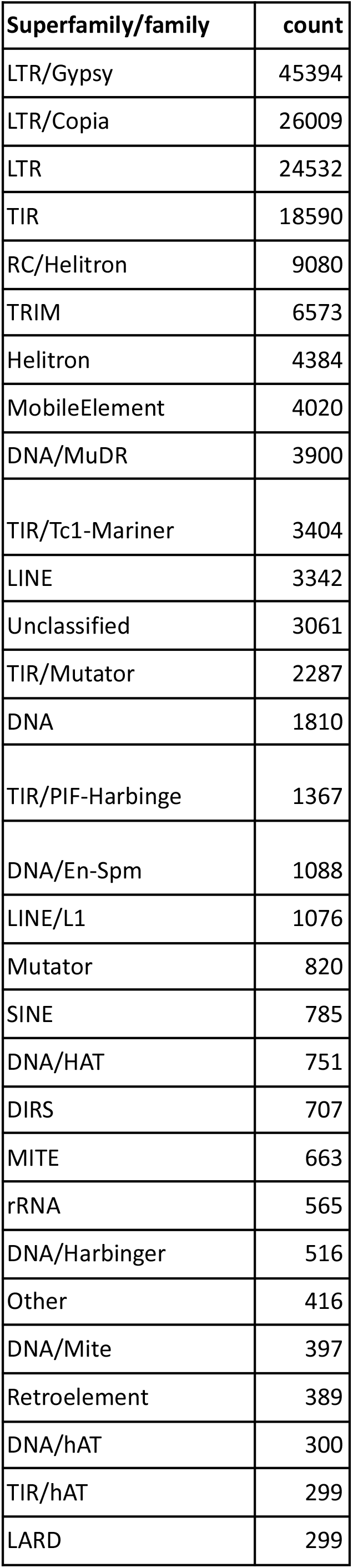
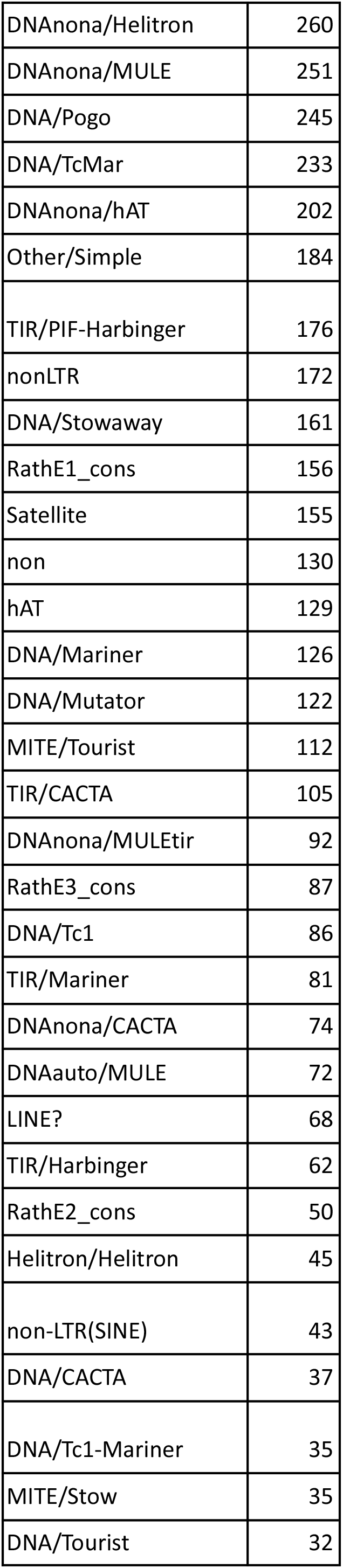
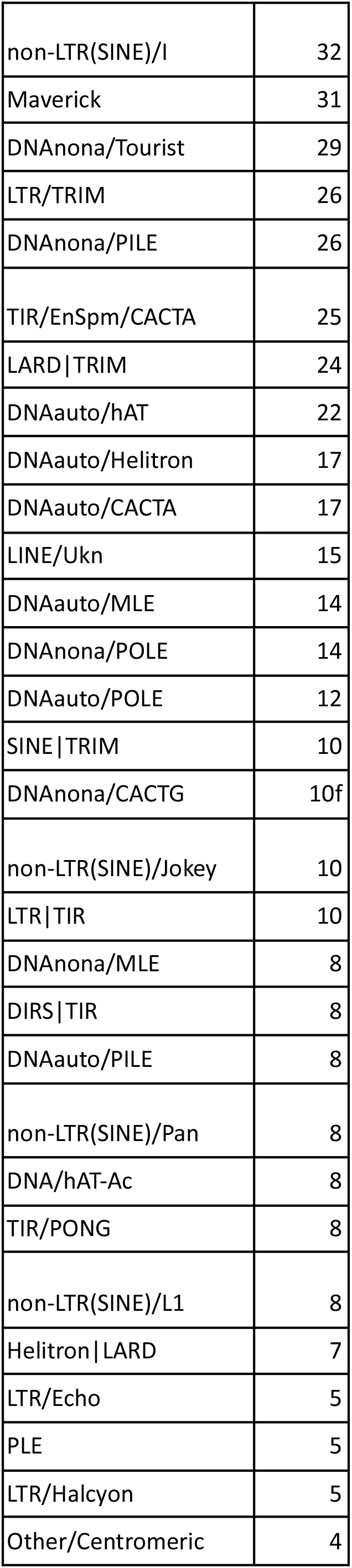
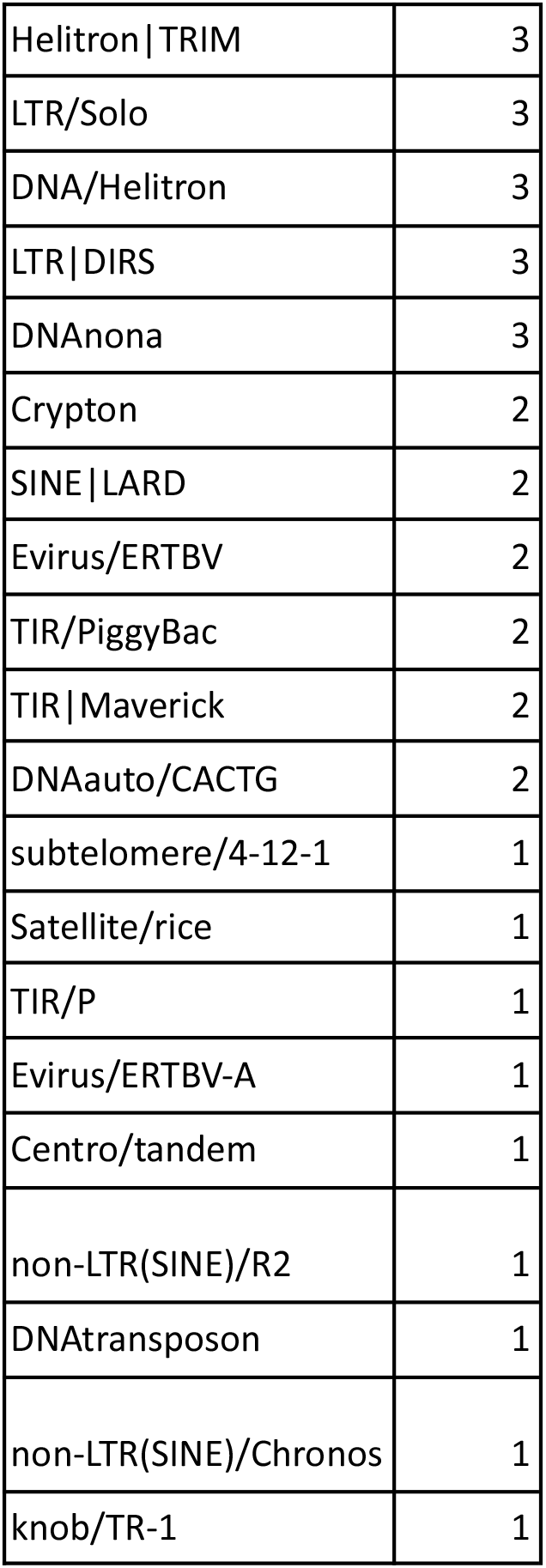
Summary of classified repeats in the nrTEplants library.

**Table S4.** Median number of repeat K-mers with 20+ copies overlapping 500bp up/downstream regions in plant genomes. Values were computed from twenty genomes from release 49 (November 2020) of Ensembl Plants annotated with Red (15) or RepeatMasker (7) with libraries REdat (20) and nrTEplants. Total k-mers are also shown.

| K | REdat<br>(n>20) | nrTEplants<br>(n>20) | Red<br>(n>20) | REdat<br>(total) | nrTEplants<br>(total) | Red<br>(total) |
| --- | --- | --- | --- | --- | --- | --- |
| 16 | 2101 | 3806.5 | 14730 | 1212499.5 | 3230821 | 6125298 |
| 21 | 1307.5 | 2285.5 | 8057 | 1178632.5 | 3150425.5 | 5925111.5 |
| 31 | 606.5 | 984 | 4256.5 | 1112352.5 | 2993877.5 | 5533979 |

**Table S5.**
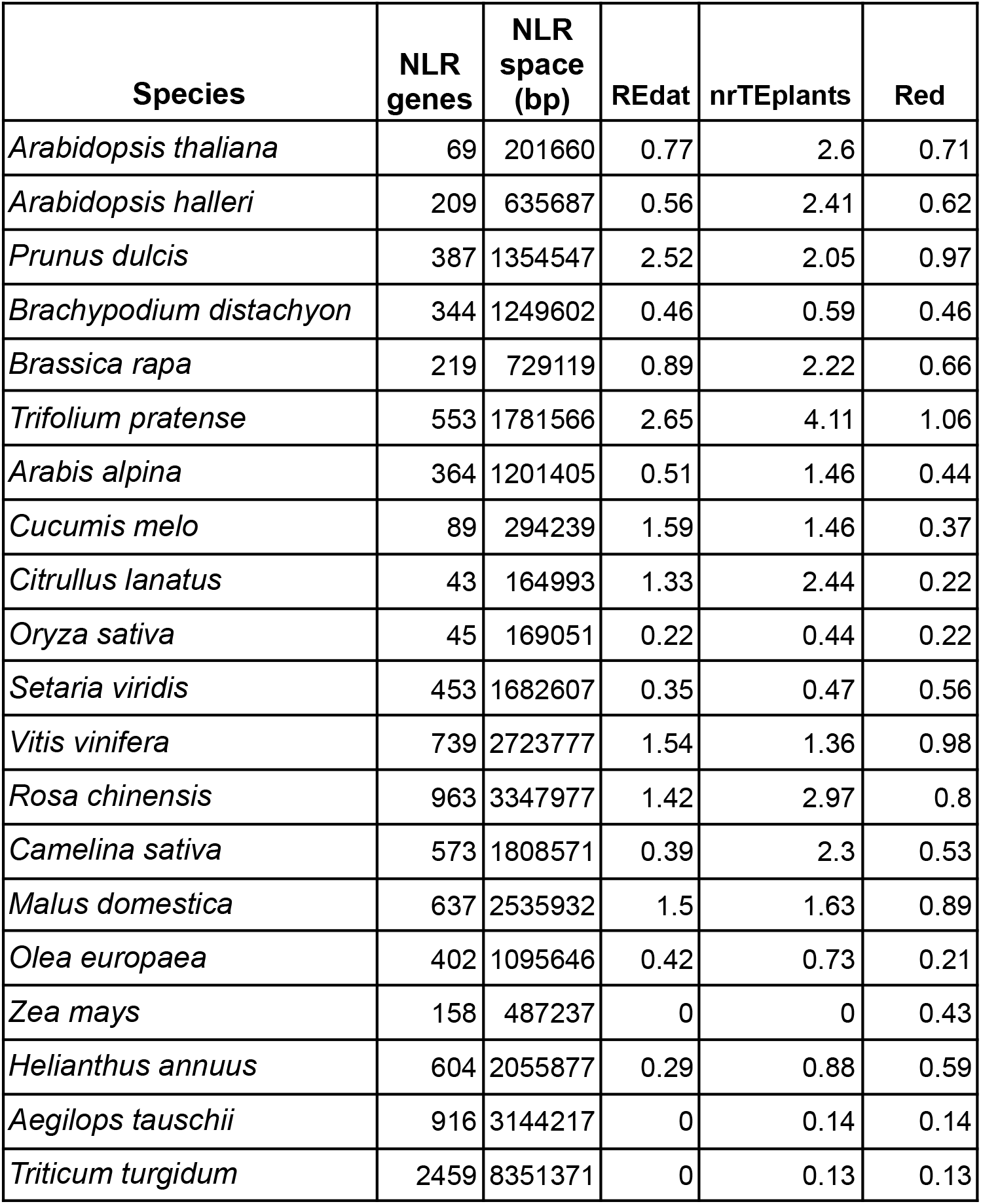
Odd ratios of NLR masking for genes overlapping > 50bp of masked sequences. Values were computed from twenty genomes from release 49 (November 2020) of Ensembl Plants annotated with Red (15) or RepeatMasker (7) with libraries REdat (20) and nrTEplants.

| Species | NLR genes | NLR space (bp) | REdat | nrTEplants | Red |
| --- | --- | --- | --- | --- | --- |
| *Arabidopsis thaliana* | 69 | 201660 | 0.77 | 2.6 | 0.71 |
| *Arabidopsis halleri* | 209 | 635687 | 0.56 | 2.41 | 0.62 |
| *Prunus dulcis* | 387 | 1354547 | 2.52 | 2.05 | 0.97 |
| *Brachypodium distachyon* | 344 | 1249602 | 0.46 | 0.59 | 0.46 |
| *Brassica rapa* | 219 | 729119 | 0.89 | 2.22 | 0.66 |
| *Trifolium pratense* | 553 | 1781566 | 2.65 | 4.11 | 1.06 |
| *Arabis alpina* | 364 | 1201405 | 0.51 | 1.46 | 0.44 |
| *Cucumis melo* | 89 | 294239 | 1.59 | 1.46 | 0.37 |
| *Citrullus lanatus* | 43 | 164993 | 1.33 | 2.44 | 0.22 |
| *Oryza sativa* | 45 | 169051 | 0.22 | 0.44 | 0.22 |
| *Setaria viridis* | 453 | 1682607 | 0.35 | 0.47 | 0.56 |
| *Vitis vinifera* | 739 | 2723777 | 1.54 | 1.36 | 0.98 |
| *Rosa chinensis* | 963 | 3347977 | 1.42 | 2.97 | 0.8 |
| *Camelina sativa* | 573 | 1808571 | 0.39 | 2.3 | 0.53 |
| *Malus domestica* | 637 | 2535932 | 1.5 | 1.63 | 0.89 |
| *Olea europaea* | 402 | 1095646 | 0.42 | 0.73 | 0.21 |
| *Zea mays* | 158 | 487237 | 0 | 0 | 0.43 |
| *Helianthus annuus* | 604 | 2055877 | 0.29 | 0.88 | 0.59 |
| *Aegilops tauschii* | 916 | 3144217 | 0 | 0.14 | 0.14 |
| *Triticum turgidum* | 2459 | 8351371 | 0 | 0.13 | 0.13 |

## Notes

https://github.com/Ensembl/plant-scripts

### Competing Interest Statement

The authors have declared no competing interest.

